# Modeling the Pancreatic Cancer Microenvironment in Search of Control Targets

**DOI:** 10.1101/2021.05.04.442611

**Authors:** Daniel Plaugher, David Murrugarra

**Affiliations:** University of Kentucky

## Abstract

Pancreatic Ductal Adenocarcinoma is among the leading causes of cancer related deaths globally due to its extreme difficulty to detect and treat. Recently, research focus has shifted to analyzing the microenvironment of pancreatic cancer to better understand its key molecular mechanisms. This microenvironment can be represented with a multi-scale model consisting of pancreatic cancer cells (PCCs), pancreatic stellate cells (PSCs), as well as cytokines and growth factors which are responsible for intercellular communication between the PCCs and PSCs. We have built a stochastic Boolean network (BN) model, validated by literature and clinical data, in which we probed for intervention strategies that force this gene regulatory network (GRN) from a diseased state to a healthy state. To do so, we implemented methods from phenotype control theory to determine a procedure for regulating specific genes within the microenvironment. We identify target genes and molecules such that the application of their control drives the GRN to the desired state by suppression (or expression) and disruption of specific signaling pathways that will eventually lead to the eradication of the cancer cells. After applying well studied control methods such as stable motifs, feedback vertex set, and computational algebra, we discovered that each produces a different set of control targets that are not necessarily minimal nor unique. Yet, we were able to gain more insight about the performance of each process and the overlap of targets discovered. Nearly every control set contains cytokines, KRas, and HER2/neu which suggests they are key players in the system’s dynamics. To that end, this model can be used to produce further insight into the complex biological system of pancreatic cancer with hopes of finding new potential targets.

## 1 Introduction

Pancreatic Ductal Adenocarcinoma is among the leading causes of cancer related deaths in the world, largely due to its aggressive nature and difficulty to detect. The pancreas is located deep in the body, which means a pancreatic tumor cannot be seen or felt during a standard doctor’s exam. What’s more, there are typically no symptoms until the cancer has spread to nearby organs. For this reason there is only a 3% five year survival rate among its victims. Symptoms can include sudden unexplained weight loss, loss of appetite, fatigue, abdominal pain, indigestion, nausea, pale stool, and dark urine [1]. Pancreatic cancer is currently the fourth highest cause of cancer related death in the United States, and it is projected to reach second by year 2030. On a global scale, pancreatic cancer is the seventh leading cause of cancer related death [2, 3].

The pancreas has two main cell types: exocrine cells and endocrine cells. Exocrine cells are responsible for digestive acids and enzymes, while endocrine cells produce hormones such as insulin and glucagon. There are many types of pancreatic cancer, but the most common is Adenocarcinoma (approximately 95% of cases) which originates in exocrine cells in the pancreatic duct lining. Islet Cell Carcinoma begins in the endocrine cells, and most cases are malignant. However, tumors that produce insulin are often benign. A functional tumor will likely produce high levels of hormones causing symptoms, whereas non-functional tumors will show less symptoms because they do not produce hormones. Other pancreatic cancer types that are more rare can include pancreaticoblastoma (mostly found in young children), pseudopapillary neoplasms (mostly found in women in their teens and 20s), ampullary cancer, adenosquamous carcinoma, squamous cell carcinoma, insulinoma, gastrinoma, and glucagonoma [1].

Therapy protocols commonly include chemotherapy, radiotherapy, and curative resection. Yet, treatments are often unsuccessful because diagnoses are frequently given at late stages of the disease, leaving little time to successfully eradicate and reverse its impact. If cancer is suspected, a diagnosis can be sought through screening, ultrasound, biopsy, computed tomography scan (CT or CAT scan), magnetic resonance imaging (MRI), percuntaneous transhepatic cholangiography (PTC), endoscopic retrograde cholangiopancreatography (ERCP), or positron emission tomography (PET). Prevention and detection can also come from being aware of known risk factors that may lead to a higher chance of cancer. These influences include gender (male), age (over 60), race (African-American), genes (BRCA1 and BRCA2 mutations), family history, smoking, chronic pancreatitis (alcoholism), obesity, diabetes, and environmental exposure to carcinogens [1].

With research focus now shifting towards analyzing the tumor microenvironment, it is becoming increasingly clear that molecules and cells surrounding the PCCs highly impact the cancer’s response to therapy. Major contributors to this environment include immune cells, endothelial cells, nerve cells, lymphocytes, dendritic cells, the extracellular matrix, and stellate cells. Recent studies have also shown that cancer cells secrete numerous types of cytokines, which play a role in intracellular communication between PCCs and PSCs [5]. It is hypothesized that a combination of these environmental factors act as a protective layer (or cocoon) for the PCCs. Therefore, it is crucial to find interventions that target the surrounding environment instead of strictly focusing on the PCCs themselves [6, 7, 8, 9, 10].

A rising field of interest is that of cancer systems biology, which often requires multidisciplinary collaboration to provide a wholistic understanding of a system’s complexities. Rather than relying strictly on traditional experimental approaches *in vivo* or *in vitro*, this field uses *in silico* models based on available data and literature. Notably, these models are often much less complex than the actual biological system due to the computational demand of high-dimensional systems. Researchers in phenotype control theory are primarily concerned with identifying key markers of the system that aid in understanding the various functions of cells and their molecular mechanisms. Therefore, models can perform simulations *in silico* to help predict outcomes and optimal targets for therapy intervention strategies. These models can also be used to simulate, validate, and optimize existing hypotheses. Recent efforts have allowed modelers to expand to multiscale models, which is an important challenge because cancer growth occurs over varying time scales, depending on the hierarchy of spatial scales. Inherently, implementing multiple scales considerably increases the model’s complexity [11, 12].

Populations of cells, multiple scales (time and size), and highly nonlinear dynamics all contribute to a significant burden for *in silico* models to overcome. There are a wide range of mathematical tools available to implement these complex models. Models are traditionally classified based on the time and population of gene products. For instance, there are techniques for continuous population with continuous time such as ordinary differential equations, discrete population with continuous time such as the Gillespie formulation, and discrete population with discrete time such as BNs, logical models, local models, and also their related stochastic counterparts [13]. Even though a multiscale model would likely provide more realistic simulations, there are currently no control methods that apply directly to multiscale models. For this reason, we use a Boolean network approximation of the multiscale model produced by Wang et al. [14] so that we can use readily available control methods.

In this project we build a BN model of pancreatic cancer, wherein we perform dynamical analysis in search of phenotype control targets. It is important to point out that here we consider a set of states from which there is no escape to be an attractor. Specifically, an attractor with a single state is called a fixed point. The long term dynamics of BNs will always converge to either a singleton fixed point or a complex attractor. With target control, it is assumed that a small subset of key nodes will be able to drive the system to a desirable state such as apoptosis [15]. We used [14] as a guide when building our model, and we were able to corroborate their results with simulations using stochastic discrete dynamical systems (SDDS) [16]. We then compare results from stable motifs (SM) [17], feedback vertex set (FVS) [18], and computational algebra (CA) [19] to analyze the control sets produced by each method. Each of these methods will be discussed in more detail in a later sections.

In Section 1 we provide the background of PCCs and motivation for the project. We also give further detail on the biological background of the regulatory network used to build our model. In Section 2, we discuss the construction of our model. Section 3 reveals the results of our analysis. Section 4 contains an overview of the methods used. In Section 5, we hold a discussion of challenges faced and potential future work, and Section 6 contains an appendix of materials such as tables, functions, and examples.

### 1.1 Network Cascades Analysis

Since we want to analyze phenotypes, we focus on specific pathways that regulate our desired end results. Wang et al. [14] provide a comprehensive analysis of the network’s cascades, and the following segment is a summary of their discoveries. The network’s topology can be found in Figure 2.

**Figure 1:**
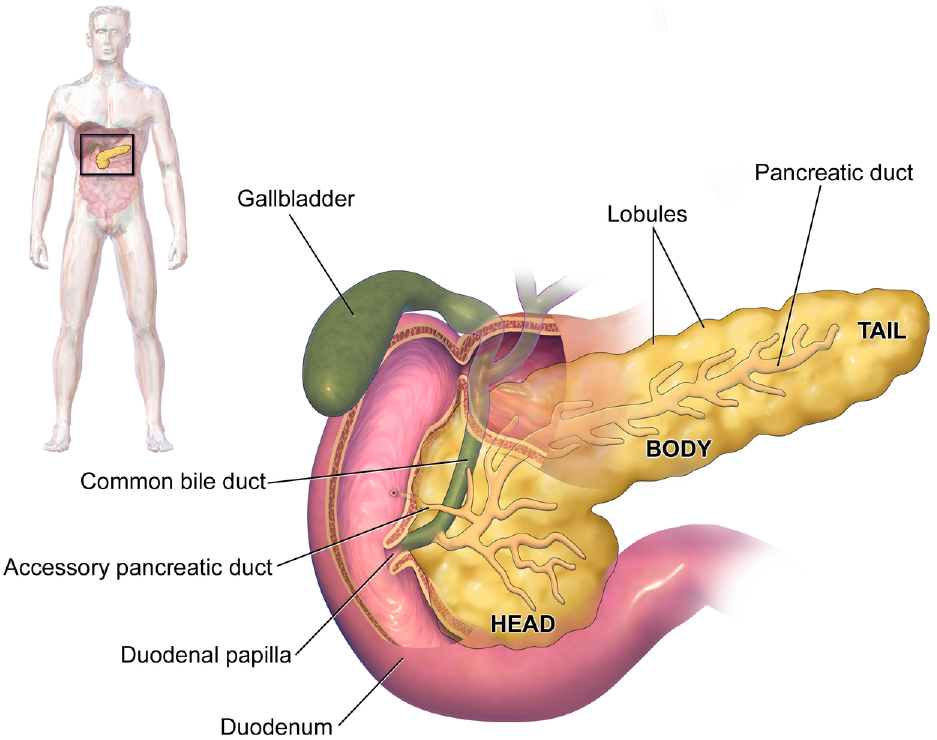
Anatomy of the Pancreas [4].

**Figure 2:**
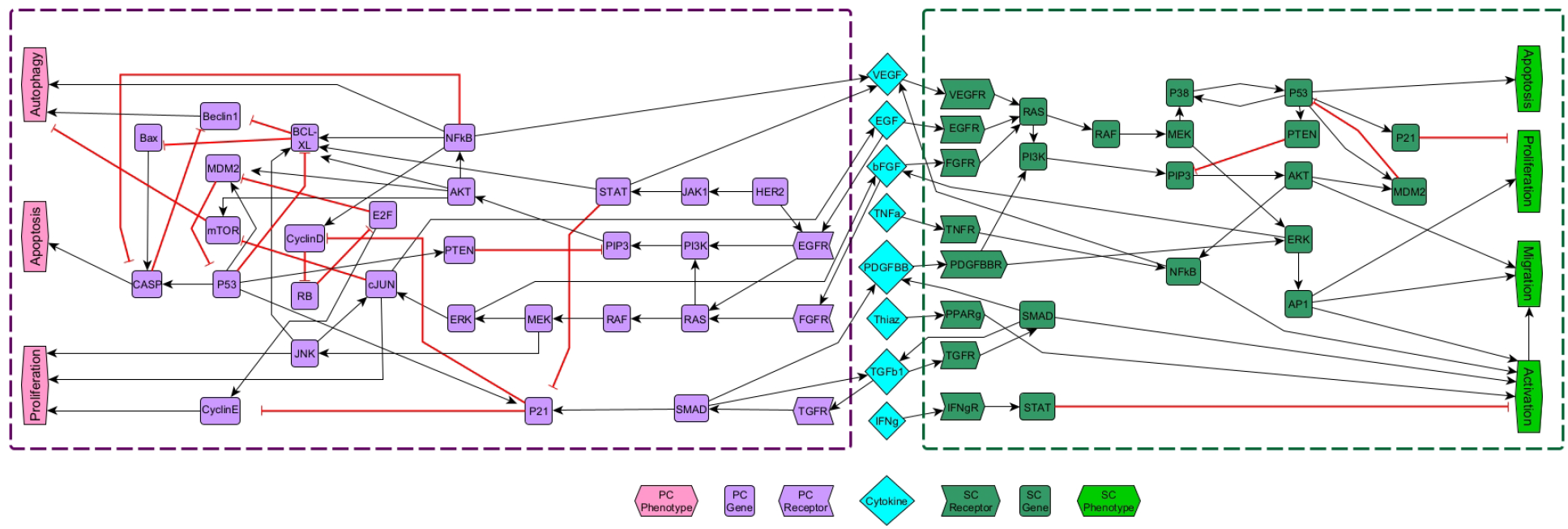
Whole gene regulatory network model of pancreatic cancer including PCCs, PSCs, and extracellular molecules. Shapes and colors of nodes indicate their function and cell type (respectively), as shown in the legend. Black barbed arrows indicate signal expression, while red bar arrows indicate suppression. The Boolean functions for each node can be found in Table 5.

In the pancreatic stellate cell, signaling pathways influencing activation of PSCs begin with PDGFBB, TGF*β*1, and TNF*α*. It was noted that the PCCs can activate the surrounding PSCs that were previously inactive by secreting these factors. Specifically, PDGFBB activates PSCs through the ERK / AP1 pathway, TGF*β*1 sustains the activation of PSCs through the TGFR / SMAD pathway, and TNF*α* prompts NF*κ*b. All these characteristics serve in promoting activation of PSCs.

Once PSCs become activated, they begin to migrate towards PCCs to form a type of protective layer. The signaling pathways influencing migration of PSCs begin with EGF, bFGF, VEGF, and PDGFBB. The growth factors will bind with their receptors and initiate migration through the MAPK / ERK pathway, also known as the RAS/ RAF /MEK /ERK pathway. Further, PDGFBB impacts migration through the PI3K / PIP3 / AKT and ERK / AP1 pathways. All these characteristics serve in promoting migration of PSCs.

Next we focus on proliferation, the rapid and uncontrollable reproduction of cells. The signaling pathways influencing proliferation in PSCs begin with EGF and bFGF, where the MAPK pathway leads to the ERK / AP1 pathway and promotes proliferation. It is noted that activated PSCs proliferate much quicker than those that are inactive. Other elements influencing proliferation are the tumor suppressers p53, p21, and PTEN which inhibit PSC proliferation.

Lastly, apoptosis is the mechanism of programmed cell death. The signaling pathways influencing apoptosis in the PSCs involve p53, which is regulated by the MAPK pathway.

Within the PCCs, there are similar regulatory pathways. Beginning with autophagy, this natural process is where cells heal themselves. The cell will break down any damaged or unnecessary components, and it will reallocate the nutrients from these processes to those that are essential. The signaling pathways influencing autophagy in PCCs involve mTOR, as well as anti-apoptotic factors NFkB and Beclin1. The PI3K / PIP3 / AKT pathway activates mTOR which inhibits autophagy, but the MEK / ERK pathway inhibits mTOR which in turn enhances autophagy. Over expression of anti-apoptotic factors such as NFkB and Beclin1 can also promote autophagy. Interestingly, it is noted that apoptosis and autophagy can inhibit each other because of indirect communication. Promoting apoptosis will inhibit autophagy, but when apoptosis is suppressed by the over expression of NFkB and Beclin1, autophagy becomes dominant and promotes PCC survival.

The signaling pathways influencing proliferation in PCCs often involve various mutations. First, the KRas oncogene mutation occurs in the precancerous stage. This then leads to continuous expression of the RAS protein, which then constantly expresses the MAPK pathway, and promotes PCC proliferation through ERK and JNK activation. Next, the HER2/neu oncogene mutation frequently occurs in the precancerous stage, where HER2 binds to EGFR and activates its cascade. Growth factors such as EGF and bFGF also promote proliferation in PCCs. It is noted that EGFR is expressed in roughly 95% of PCCs and promotes proliferation through the RAS / RAF / MEK / JNK pathway. Another option is the activation of the MAPK / cJUN pathway which leads to the secretion of more EGF molecules that then bind and over express EGFR to further promote proliferation. Similarly, bFGF can also promote proliferation through both MAPK and RAF / MEK / JNK pathways. As before, the MAPK pathway can lead to the secretion of more bFGF molecules that then bind and over express FGFR to further promote proliferation. All of these aspects assist in the promotion of proliferation in PCCs.

The signaling pathways influencing apoptosis in PCCs begin with TGF*β*1, and mutated oncogenes. In the early development of PCCs, TGF*β*1 will activate SMADs which then relocate to the nucleus to begin apoptosis. However, just as the mutated oncogenes KRas and HER2/neu promote proliferation, they also inhibit apoptosis by downregulating caspases (CASP) through the PI3K / AKT / NF*κ*B pathway and by inhibiting Bax through the PI3K / PIP3 / … / Bax pathway.

Finally, we address the intercellular communication between PSCs and PCCs which is conducted through growth factors and cytokines. Typically, PSCs exist quiescently in the periacinar, perivascular, and periductal space. But diseased PSCs will be activated by growth factors, cytokines, and oxidant stress secreted or induced by PCCs as mentioned above. Once activated, PSCs begin rapidly proliferating and migrating while also heavily secreting cytokines and molecules that adhere to surrounding cells. Likewise, the PSCs promote PCCs by protecting them from various therapy protocols. This is achieved by the PSCs’ secretion of stromal-derived factor 1, bFGF, secreted protein acidic and rich in cysteine, matrix metalloproteinases, small leucinerich proteoglycans, periostin and collagen type I that mediate effects on tumor growth, invasion, and metastasis [14].

## 2 Building the model

In [14], a rule-based model was presented with BioNetGen and StatMC using a combination of discrete and continuous rules. They constructed a multi-cellular model of the pancreatic cancer system microenvironment shown in Figure 2. Notice that the figure shows the signaling pathways for the PCCs, PSCs, and molecules such as growth factors and cytokines that are the ligands for the receptors within the PCCs and PSCs. Using statistical model checking, they were able to test ten different properties with three different scenarios of target control strategies.

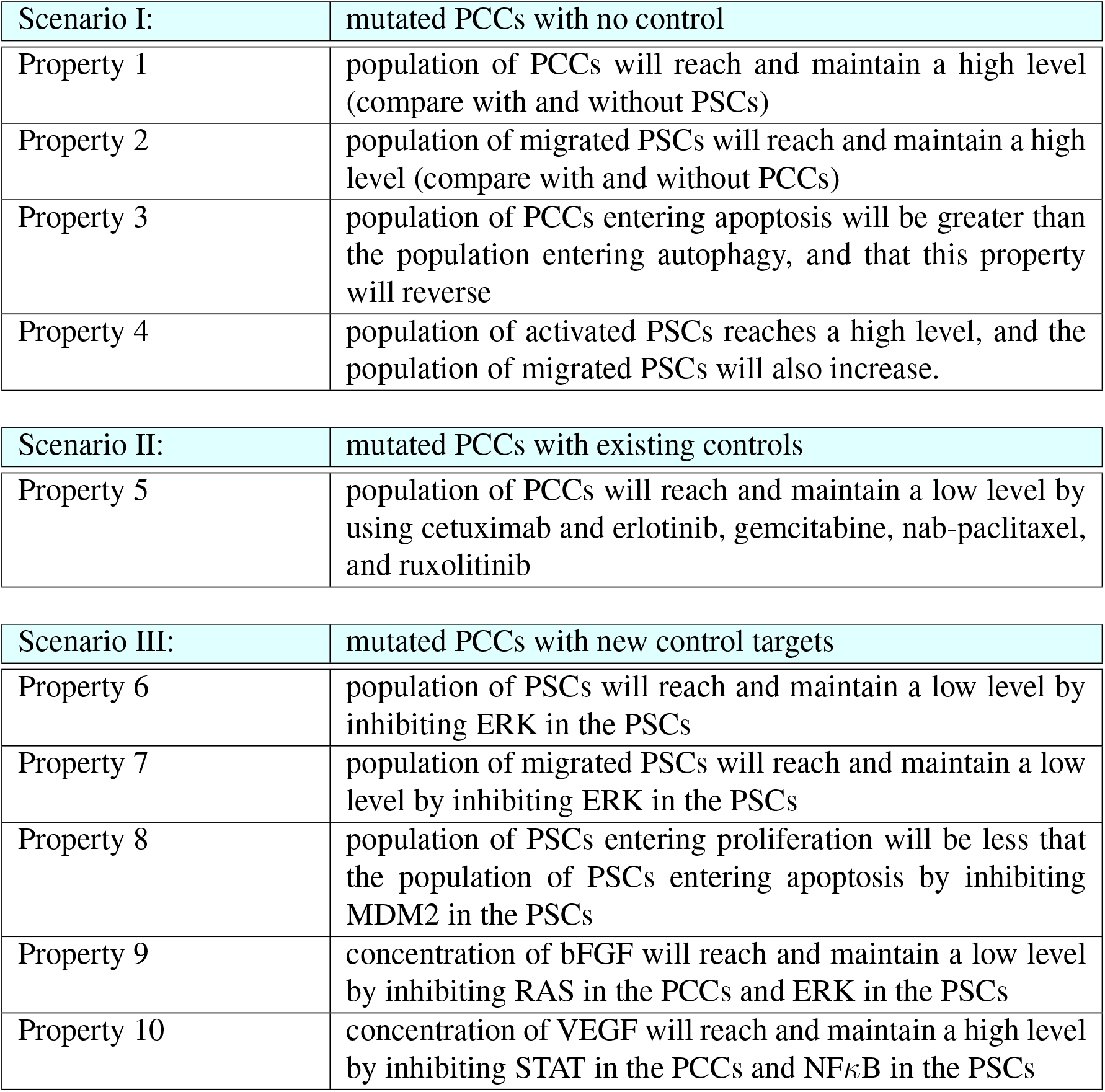

Downstream signaling and information transformation was encoded using formulas in a Boolean network, where each node is either ON or OFF (1 or 0). In other words, the node is either being expressed or suppressed. We used Properties 1-10 to write the functions and fine tune our simulator, which uses the SDDS framework described in Section 4. Each function indicates the next state of the given node in terms of the current states of that nodes’ regulators and is justified via literature (see Table 5). We then determine the state of the entire system by the state of each node at current time, noting phenotypic markers. Node expression is written as OR statements, while suppression is written as AND NOT. The exceptions to this rule are PCC proliferation, PSC migration, and PSC activation because of their upstream signaling. In Figure 2, the shapes and colors of nodes indicate their function and cell type, respectively. Black barbed arrows indicate signal activation, while red bar arrows indicate deactivation.

For an example of function construction, PSC RAS has multiple inputs including VEGFR, EGFR, and FGFR. Since all inputs are promoting RAS, as seen in Figure 2, our function is

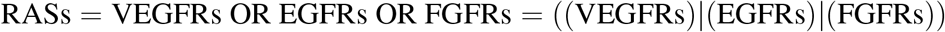

Similarly, notice that PSC p53 is inhibited by MDM2 but promoted by p38. Therefore, our function is

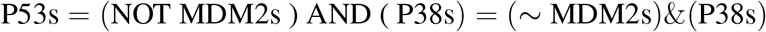

Once we developed a working model, we were able to begin analyzing the system with the various methods mentioned previously. For calibration purposes, we split our model into two separate systems while keeping the cytokines intact. Splitting our model into separate PCC and PSC systems allowed us to verify that our dynamics aligned with the findings in literature. It is also important to note that angiogenesis was excluded for computational efficiency, but it can easily be calculated using the VEGF function value.

## 3 Results

Using our model, we sought to identify targets such that their control drives the cancer system into a healthy state by suppression (or expression) and disruption of specific signaling pathways. After applying well studied control methods such as stable motifs, feedback vertex set, and computational algebra, we discovered that each produces a different set of control targets that are not necessarily minimal nor unique. The aforementioned computational expense of the large system was too great to apply all these methods directly. To accommodate, we applied the control methods to a reduced network, and then we confirmed that the controls hold for the whole system using our simulator. All computations were performed on an Intel(R) Core i3-4170T CUP @3.2GHz with 12 GB of RAM and a 64-bit operating system.

### 3.1 Model Reduction

In a BN, the magnitude of the state space for *n* genes is 2^*n*^. So we naturally see that an increase of GRN size will exponentially increase the computational burden for its analysis, and therefore, brute-force methods for small systems are not sufficient. Consequently, we implemented well studied reduction techniques to reduce our system from sixty-nine nodes to twenty-two nodes, which is a 68% reduction (see Figure 3). First we removed nodes with one input and one output, but we maintained nodes with self-loops and phenotypes as biomarkers [20]. When a node was deleted, its function values were substituted directly into its downstream signal recipient(s). The only exception to the phenotype retention rule was cancer cell apoptosis in order to aid in computation efficiency, where we instead use CASP as its marker. Next, we removed nodes with either one input and multiple outputs, or vice versa. Lastly, we removed nodes with low connectivity relative to the remaining nodes. These techniques have been shown to preserve fixed points but not complex attractors, yet, our results indicate a conservation of attractors [21, 22].

**Figure 3:**
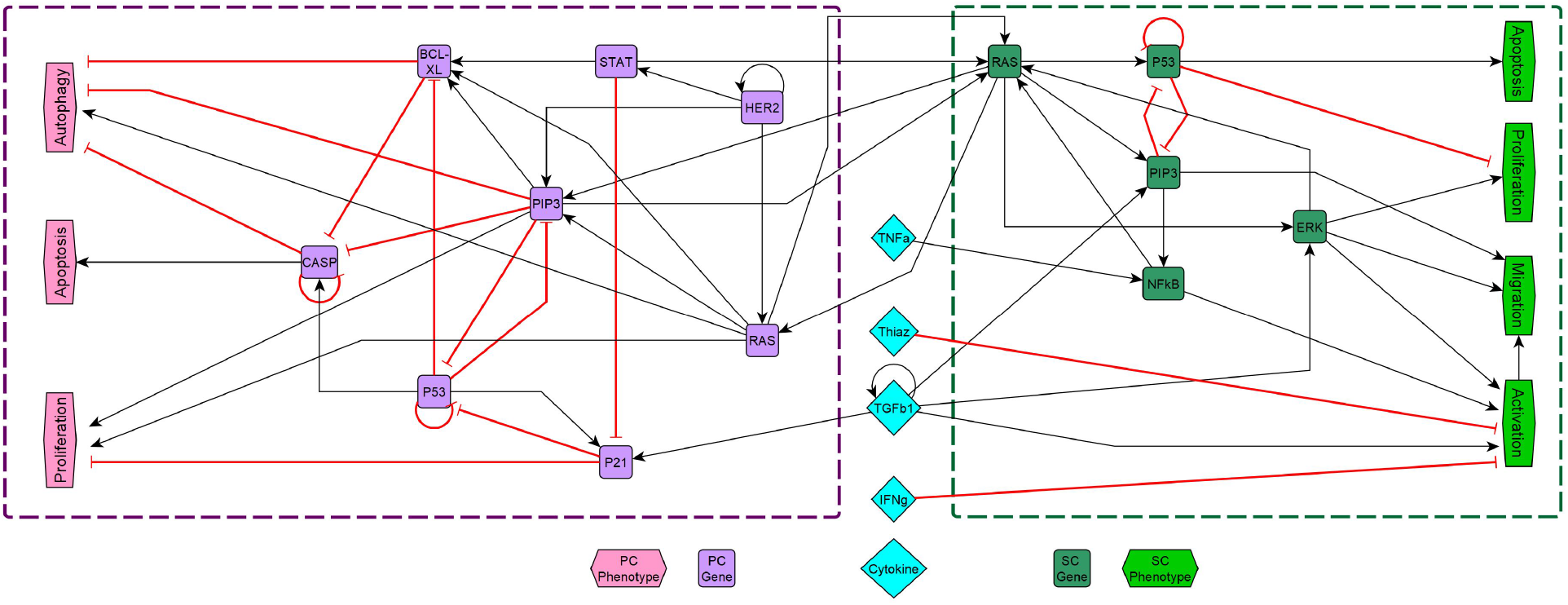
Reduced gene regulatory network model of pancreatic cancer including PCCs, PSCs, and extracellular molecules. The reduced model contains twenty-two nodes, compared to the whole model with sixty-nine nodes. As before, the shapes and colors of nodes indicate their function and cell type (respectively), as shown in the legend. Black barbed arrows indicate signal expression, while red bar arrows indicate suppression. The Boolean functions for each node can be found in Table 6. For examples of reductions, see Figures 4–9 in the Appendix.

### 3.2 Long-Term Dynamics

Publicly available software utilizing SMs [17, 23] revealed that there are a total of thirty-six attractors, eight of which are fixed points (see Table 8). Once all attractors were identified, we were able to use our simulator to confirm that the model always converged to an attractor given sufficient run time. Typically fifty time steps within our simulator was more than enough. We then applied the SM code to the reduced model in Figure 3 and again arrived at the same thirty-six attractors (see Table 7), revealing that attractors were maintained. As such, we believe there is an underlying mechanism within our reductions that preserves attractors and remains to be explored.

### 3.3 Phenotype Control

In this section we describe our results from applying three methods for phenotype control: Stable Motifs (SM), Feedback Vertex Set (FVS), and Computational Algebra (CA).

#### 3.3.1 Stable Motifs

In Table 1, we list the control sets discovered with SMs for each of the four attractors associated with cancer cell apoptosis. We draw attention to the fact that each control set contains the cytokines, HER2 and cancer cell RAS. Note that the amount of sets is a result of the networks topology, with combinations of the source nodes Thiazolidinedione and IFN*γ* (see Figure 2). We interpret these results to mean that their expressions (or supressions) are not necessary for controlling the system under the SM structure. Thus, the smallest set of controls to achieve PCC apoptosis with stable motifs would contain five nodes (TNF*α*, TGF*β*1, HER2p, RASp, RASs) and all are OFF. We subsequently used these targets in our simulator to confirm arrival to the desired attractors.

**Table 1:** Set of control targets associated with PCC apoptosis. These control sets were generated using SMs applied to the reduced network. We correspond OFF with the Boolean identifier of 0 in red, and ON with 1 in green. There are four apoptotic PCC attractors (A, B, C, D -see Table 8), each with three sets of targets.

| Apoptosis |  |  |  |  |  |
| --- | --- | --- | --- | --- | --- |
| A |  |  | B |  |  |
| $TNF\alpha=0$ | $TNF\alpha=0$ | $TNF\alpha=0$ | $TNF\alpha=0$ | $TNF\alpha=0$ | $TNF\alpha=0$ |
| Thiaz.=0 | Thiaz.=0 | Thiaz.=0 | Thiaz.=1 | Thiaz.=1 | Thiaz.=1 |
| $TGF\beta1=0$ | $TGF\beta1=0$ | $TGF\beta1=0$ | $TGF\beta1=0$ | $TGF\beta1=0$ | $TGF\beta1=0$ |
| $IFN\gamma=0$ | $IFN\gamma=0$ | $IFN\gamma=0$ | $IFN\gamma=0$ | $IFN\gamma=0$ | $IFN\gamma=0$ |
| HER2p=0 | HER2p=0 | HER2p=0 | HER2p=0 | HER2p=0 | HER2p=0 |
| RASp=0 | RASp=0 | RASp=0 | RASp=0 | RASp=0 | RASp=0 |
| RASs=0 | $NF\kappa Bs=0$ | $PIP3s=0$ | RASs=0 | $NF\kappa Bs=0$ | $PIP3s=0$ |
|  | ERKs=0 | ERKs=0 |  | ERKs=0 | ERKs=0 |
| C |  |  | D |  |  |
| $TNF\alpha=0$ | $TNF\alpha=0$ | $TNF\alpha=0$ | $TNF\alpha=0$ | $TNF\alpha=0$ | $TNF\alpha=0$ |
| Thiaz.=0 | Thiaz.=0 | Thiaz.=0 | Thiaz.=1 | Thiaz.=1 | Thiaz.=1 |
| $TGF\beta1=0$ | $TGF\beta1=0$ | $TGF\beta1=0$ | $TGF\beta1=0$ | $TGF\beta1=0$ | $TGF\beta1=0$ |
| $IFN\gamma=1$ | $IFN\gamma=1$ | $IFN\gamma=1$ | $IFN\gamma=1$ | $IFN\gamma=1$ | $IFN\gamma=1$ |
| HER2p=0 | HER2p=0 | HER2p=0 | HER2p=0 | HER2p=0 | HER2p=0 |
| RASp=0 | RASp=0 | RASp=0 | RASp=0 | RASp=0 | RASp=0 |
| RASs=0 | $NF\kappa Bs=0$ | $PIP3s=0$ | RASs=0 | $NF\kappa Bs=0$ | $PIP3s=0$ |
|  | ERKs=0 | ERKs=0 |  | ERKs=0 | ERKs=0 |

#### 3.3.2 Feedback Vertex Set

Using publicly available software [18, 24], we were able analyze both the reduced and whole systems when searching for FVSs. This particular software approximates the minimum FVS of a directed graph [18]. Once a FVS is identified, we can use previously established attractor information to determine the node values of those in the desired attractors.

In Tables 2 and 3, we list the control sets for the four attractors associated with cancer cell apoptosis within both the reduced and whole models. As before, each set contains the cytokines, HER2, and cancer cell RAS. We again recognize that the combinations of Thiazolidinedione and IFN*γ* produce four total sets, and cancer cell p53 is designated as a cycle, which means they are not necessary for control under the FVS framework. Thus, the smallest set of controls discovered with FVS would contain six nodes (TNF*α*, TGF*β*1, p53s, RASs, HER2p, RASp) and all are OFF. The whole model added cancer cell MEK and Beclin, but the nodes for the reduced model still hold for the whole model because the smallest set discovered is a subset of the larger. We confirmed that these results always force convergence to the correct attractors given sufficient run time. In Section 5, it is discussed that the FVS algorithm does not produce an exhaustive list, but rather a minimal set. So it is possible to get varying sets of controls, however, the magnitudes will remain the same.

**Table 2:** Set of control targets associated with PCC apoptosis in the reduced network.

| Apoptosis |  |  |  |  |
| --- | --- | --- | --- | --- |
| TNFα | 0 | 0 | 0 | 0 |
| Thiaz. | 0 | 1 | 0 | 1 |
| TGFβ1 | 0 | 0 | 0 | 0 |
| IFNγ | 0 | 0 | 1 | 1 |
| RASs | 0 | 0 | 0 | 0 |
| P53s | 0 | 0 | 0 | 0 |
| HER2p | 0 | 0 | 0 | 0 |
| RASp | 0 | 0 | 0 | 0 |
| P53p | X | X | X | X |

**Table 3:** Set of control targets associated with PCC apoptosis in the whole network.

| Apoptosis |  |  |  |  |
| --- | --- | --- | --- | --- |
| TNFα | 0 | 0 | 0 | 0 |
| Thiaz | 0 | 1 | 0 | 1 |
| TGFβ1 | 0 | 0 | 0 | 0 |
| IFNγ | 0 | 0 | 1 | 1 |
| RASs | 0 | 0 | 0 | 0 |
| P53s | 0 | 0 | 0 | 0 |
| HER2p | 0 | 0 | 0 | 0 |
| MEKp | 0 | 0 | 0 | 0 |
| Beclinp | 0 | 0 | 0 | 0 |
| P53p | X | X | X | X |

#### 3.3.3 Computational Algebra

For the algebraic control approach, we used the polynomial form of the Boolean functions seen in Table 6 to implement the method described in [19] which includes node and edge controls. For node control, we encode the nodes of interest as control variables, and then the control objective is described as a system of polynomial equations that is solved by computational algebra techniques.

In Table 4, we list the control sets for node control of apoptosis within the reduced model with the objective to block PCC proliferation. CA provided sets that had no overlap with the other methods used. Whats more, the magnitudes of the sets are significantly smaller comparatively. Here we see sets of either of singleton nodes or pairs targets. We confirmed that these results always force convergence to a healthy state given sufficient run time, but new attractors had been created. This phenomenon is discussed further in the Methods section.

**Table 4:** Target sets from CA node control. We indicate expression with ON in green and suppression with OFF in red. These sets are either singleton controls or pairs.

|  |
| --- |
| p53p=ON |
| RASp=OFF |
| PIP3p=OFF |
| ERKs=OFF RASp=OFF |
| ERKs=OFF PIP3p=OFF |
| RASp=OFF BCLXLp=OFF |

## 4 Methods

### 4.1 Stochastic Boolean Networks

Discrete models of GRNs can involve stochastic processes depending on the update schedules chosen. Synchronous update schedules produce deterministic dynamics, wherein nodes are all updated simultaneously. On the other hand, asynchronous update schedules produce stochastic dynamics, wherein a randomly selected node is updated at each time step. The SDDS framework (developed in [13]) incorporates Markov chain tools to study long-term dynamics of Boolean networks. SDDS uses parameters based on designated propensities to model node (and pathway) signal activation and deactivation, also referred to as degradation. More precisely, an SDDS of the variables (*x*_1_, *x*_2_, …, *x*_*n*_) is a collection of *n* triples

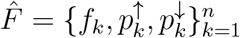

where for *k* = 1, …, *n*,

- *f*_*k*_ : {0, 1}^*n*^→ {0, 1} is the update function for *x*_*k*_
- 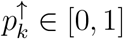 is the activation propensity
- 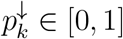 is the deactivation propensity

Here, the parameters 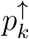 and 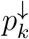 introduce stochasticity. For example, an activation of *x*_*k*_(*t*) at the next time step (i.e. *x*_*k*_(*t*) = 0, *f*_*k*_(*x*_1_(*t*), …, *x*_*n*_(*t*)) = 1, and *x*_*k*_(*t* + 1) = 1) occurs with probability 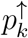. An SDDS can be represented as a Markov Chain via its transition matrix, which can be viewed as transition probabilities between various states of the network. Elements of the transition matrix *A* are determined as follows: consider the set *S* = {0, 1}^*n*^ consisting of all possible states of the network. Suppose *x* = (*x*_1_, …, *x*_*n*_) ∈ *S* and *y* = (*y*_1_, …, *y*_*n*_) ∈ *S*. Then, the probability of transitioning from *x* to *y* is

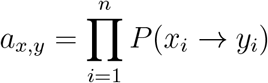

where

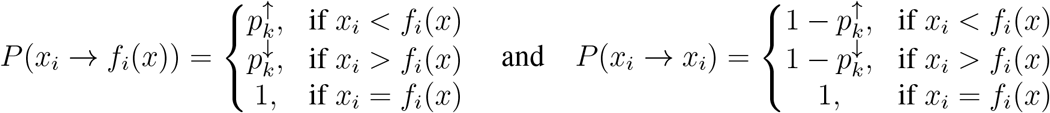

It follows that *P* (*x*_*i*_ → *y*_*i*_) = 0 for any *y*_*i*_ *∉* {*x*_*i*_, *f*_*i*_(*x*)}. Therefore, we achieve *A* = [*a*_*x,y*_]_*x,y*∈*S*_. It is important to note that this framework preserves fixed points of the system, regardless of performing synchronous or asynchronous updates. Although the basins of attraction will change due to stochasticity, other features such as limit cycles and strongly connected components are not necessarily maintained [13]. It is also noted that when propensities are set to *p* = 1, we have a traditional BN. While SDDS does not provide insight for control targets, it is the foundation of our simulator.

### 4.2 Stable Motifs

The impact of numerous regulators on a single node can be addressed and analyzed with the method of stable motifs [17]. By definition, a stable motif is a strongly connected subgraph of the expanded graph that satisfies the following:

1. it contains either a node or its complement but not both
2. it contains all inputs of its composite nodes (if any exist)

First, implement the expanded network that is used to add information about the combinatorial interaction and signs of nodes. Composite nodes represent the AND interaction and complementary nodes represent the NOT interaction. Each original node *i* is denoted by *x*_*i*_ in the expanded graph, and a complementary node (∼ *x*_*i*_) is added if the original node represented suppression. Then, all NOT functions are replaced by its appropriate complementary node in the function. Next, edges are included where each edge is a positive regulation, contrary to the original wiring diagram [15]. The second step is to make distinctions between OR rules and AND rules by using composite nodes for each function including ANDs. To do this, the functions must be in disjunctive normal form in order to uniquely determine edges. A special node is included for AND rules, and edges are drawn from the non-composite nodes of the network that form the actual composite rule. It is noted that the benefit of such an action is that the reader is able to see all regulatory functions simply from the topology of the expanded network. Now that the expanded graph is complete, using the definition above we can search for SMs within the network. The group of nodes included in the SM represent partial fixed points, from which the remaining nodes can be calculated using the original Boolean functions [15].

Zañudo and Albert implemented SMs in a Java library, using the algorithms and reduction techniques described in [17, 23]. We used their program (https://github.com/jgtz/StableMotifs) to find our attractors and control targets.

### 4.3 Feedback Vertex Set

Recall that a tree is an undirected graph in which any two vertices are connected by exactly one path, that is, a connected acyclic undirected graph. A forest is defined as an undirected graph in which any two vertices are connected by at most one path, that is, an acyclic undirected graph, or a disjoint union of trees [25]. By definition, a FVS of a graph is a minimal set of nodes whose removal leaves the graph without cycles. Formally, define a graph *G* = (*V, E*) that consists of a finite set of vertices *V* (*G*) and a set of edges *E*(*G*). Then a FVS of *G* is a subset of vertices *V* ′ ⊆ *V* (*G*) such that the removal of *V*′ from *G*, along with all edges incident to *V′*, results in a forest [26]. FVS control uses the nodes in a FVS for network control [27, 28] and has been successfully applied to a variety of networks [18].

Algorithms in [18, 24] were use to develop a python software to find FVSs. We used their program (https://github.com/jgtz/FVS_python3) to find control targets for our system.

### 4.4 Computational Algebra

The method based on computational algebra described in [19] seeks two types of controls: nodes and edges. These can be achieved biologically by blocking effects of the products of genes associated with nodes, or targeting specific gene communications. Let the function 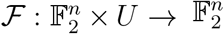 denote a Boolean network with control, where *U* is a set of all possible controls. Then, for some *u* ∈*U*, the new system dynamics are *x*(*t* + 1) = ℱ (*x*(*t*), *u*). That is, each coordinate *u*_*i,j*_ ∈ *u* encodes the control of edges as follows: consider the edge from nodes *x*_*i*_ → *x*_*j*_ in a given diagram. It follows by construction that

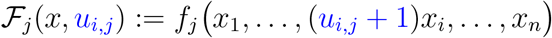

which gives

- Inactive control:

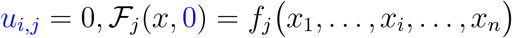
- Active control (edge deletion):

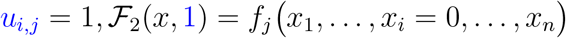

The definition of edge control can therefore be applied to many edges, obtaining 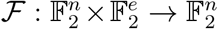 where *e* is the number of edges in the diagram.

Next, we consider control of node *x*_*i*_ from a given diagram. By construction

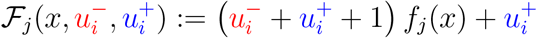

which yields

- Inactive control:

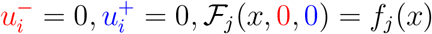
- Node *x*_*i*_ deletion:

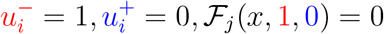
- Node *x*_*i*_ expression:

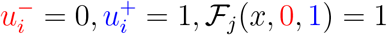
- Negated function value (irrelevant for control):

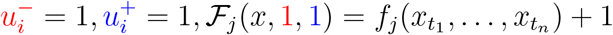

Using these definitions, we can achieve three types of objectives. Let *F* = (*f*_1_, …, *f*_*n*_) : 𝔽^*n*^ → 𝔽^*n*^ where 𝔽 = {0, 1} and *µ* = {*µ*_1_, …, *µ*_*n*_} is a set of controls. Then we can:

- *Generate new attractors*. If *y* is a desirable state (ex. apoptosis), but it is not currently an attractor, we find a set *µ* so that we can solve

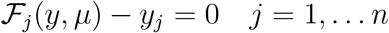
- *Block transitions or remove attractors*. If *y* is an undesirable attractor (ex. proliferation), we want to find a set *µ* so that ℱ (*y, µ*) ≠ *y*. In general, we can use this framework to avoid transitions between states (say *y* → *z*) so that ℱ (*y, µ*) ≠ *z*. So we can solve

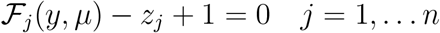
- *Block regions*. If a particular value of *y*, say *y*_*k*_ = *a*, triggers an undesirable pathway, then we need all attractors to satisfy *y*_*k*_ ≠ *a*. So we find a set *µ* so that the following system has no solution

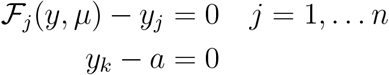

Notably, the Boolean functions *F* must be written as polynomials. To complete the control search we then compute the Grobner basis of the ideal associated with the given objective. For example, if we generate new attractors, we find the Grobner basis for

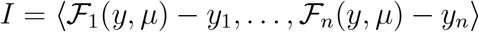

Therefore, we can determine all controls that solve the system of equations and detect combinatorial actions for the given model [19]. We used the software Macaulay2 [29] to find the fixed points and controls for our system.

## 5 Discussion and Conclusion

In this project, we developed a Boolean model of pancreatic cancer that captures properties found in literature [14]. These various traits include examining the impact and interaction of PSCs with PCCs, the impact of activated PSCs, the efficacy of known controls, and the efficacy of new controls. We were then able to confirm thirty-six total attractors, which represent phenotypic states of the system. These have been categorized into representatives for PCC proliferation, PCC autophagy, and PCC apoptosis in Tables 7 and 8. To eradicate PCCs, we focused on finding targets to promote cancer cell apoptosis or block PCC proliferation.

However, the size of the original system lead to high computational demand. To account for this issue, we used model reduction techniques [20, 21, 22] that preserve long term system dynamics. After the model was reduced from sixty-nine nodes to twenty-two, we applied stable motifs, feedback vertex set, and computational algebra to find appropriate control targets. Importantly, all controls of the reduced model were confirmed to hold in the original model via simulation. We also discovered that cytokines were key players in the downstream signaling process. Nearly every control strategy involved these extra-cellular molecules, even though some are more essential than others. Along those same lines, RAS and HER2 proved to be important across most strategies, which was confirmation of the cascade analysis in Section 1.

Through stable motifs, we were able to gain insight into the amount of attractors as well as potential control targets. However, the size of the whole model was too taxing for the software used. Though termination was not achieved, we still retrieved critical attractor information along the way. After using the reduced model, we confirmed that the attractors were maintained and discovered control sets that hold for both the reduced and whole systems. Notably, the sets produced do not appear to be minimal due to the combinations of source nodes Thiazolidinedione and IFN*γ*.

We note that a FVS is not unique for a given graph and that the problem of finding a FVS is NP-complete [30]. By definition, a FVS is a minimal set of nodes whose removal makes the system acyclic. This means all sets produced will have the same magnitude. Fortunately, there is available software to approximate a minimal FVS. We discovered that different sets were expelled when applied to the whole model, yet the reduced model did not reveal such information. The following are alternate sets found for the whole model:

- {Beclin1p, TGF*β*1, HER2p, P53s, Thiaz, MDM2p, RASs, RAFp, TNF*α*, IFN*γ*}
- {P53p, TGF*β*1, HER2p, P53s, Thiaz, Beclin1p, RASs, RAFp, TNF*α*, IFN*γ*}
- {Beclin1p, TGF*β*1, HER2p, P53s, Thiaz, MDM2p, MEKp, RASs, TNF*α*, IFN*γ*}
- {Beclin1p, TGF*β*1, HER2p, P53s, Thiaz, MDM2p, RASs, RASp, TNF*α*, IFN*γ*}
- {P53p, TGF*β*1, HER2p, P53s, Thiaz, CASPp, RASs, RAFp, TNF*α*, IFN*γ*}

As before, we noted that the sets discovered could be further reduced due to the combinations of source nodes Thiazolidinedione and IFN*γ*, as well as oscillating oscillating nodes.

Computational algebra produced much smaller sets while providing an alternative to other control methods. As within the other methods, we applied CA to the reduced model and confirmed the results hold for the whole model. A feature of the CA method is the possible creation of new attractors.

Finally, it is clear that the methods applied in this paper might produce a large number of control strategies. However, many of these may not be biologically feasible due to the size of the set or the controllability of the node. For example, we have drawn attention to source nodes that appear nonessential, and it is known that p53 is not a feasible control. It remains to find a strategy for ranking these sets according to some scoring function. We also know mutations play a major role in system dynamics, and we intend to further study their impact using clinical data.

## Acknowledgements

DM thanks Boris Aguilar for the discussions and suggestions during in the initial stage of this project.

## 6 Appendix

### Functions

**Table 5:**
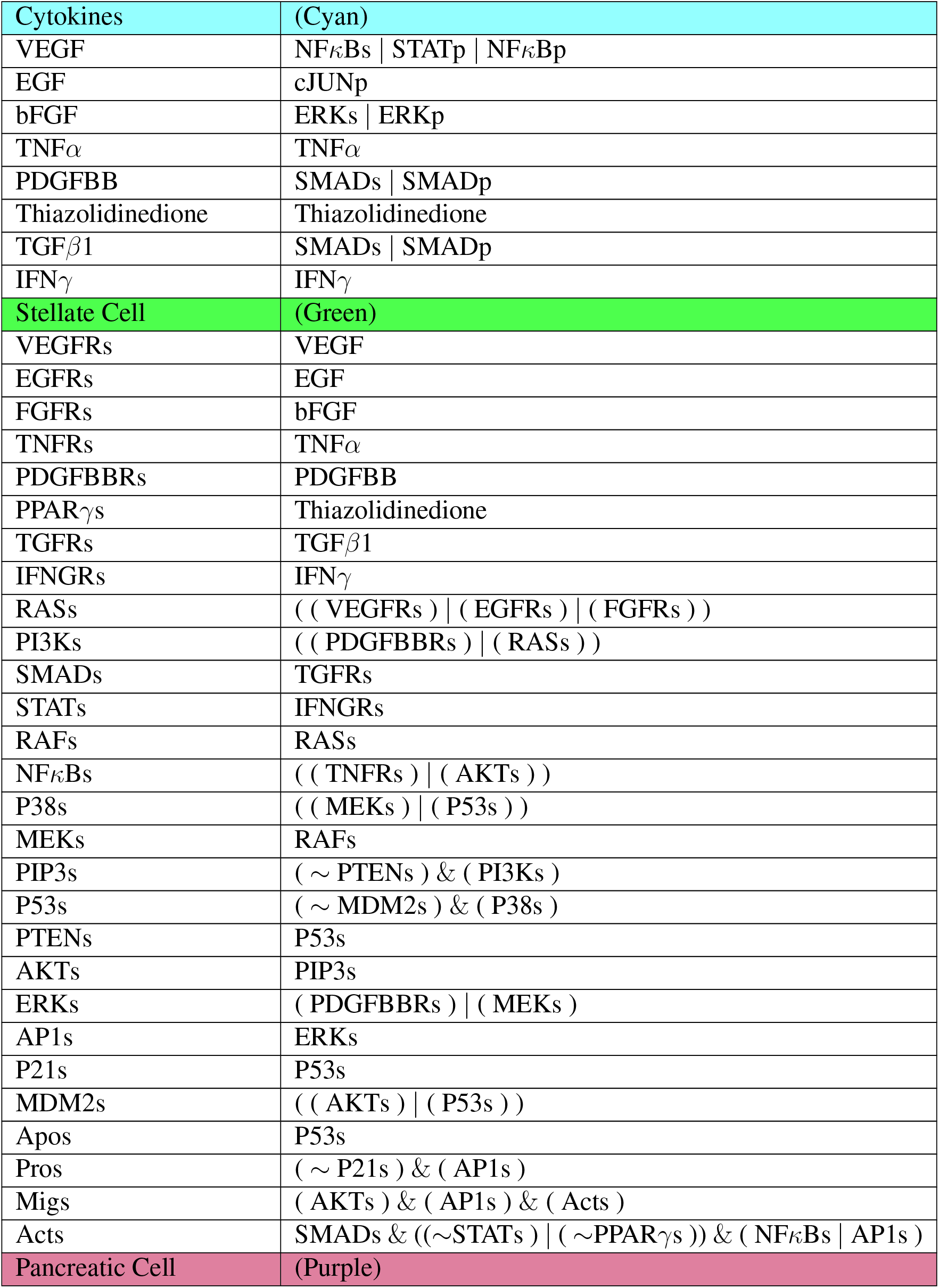
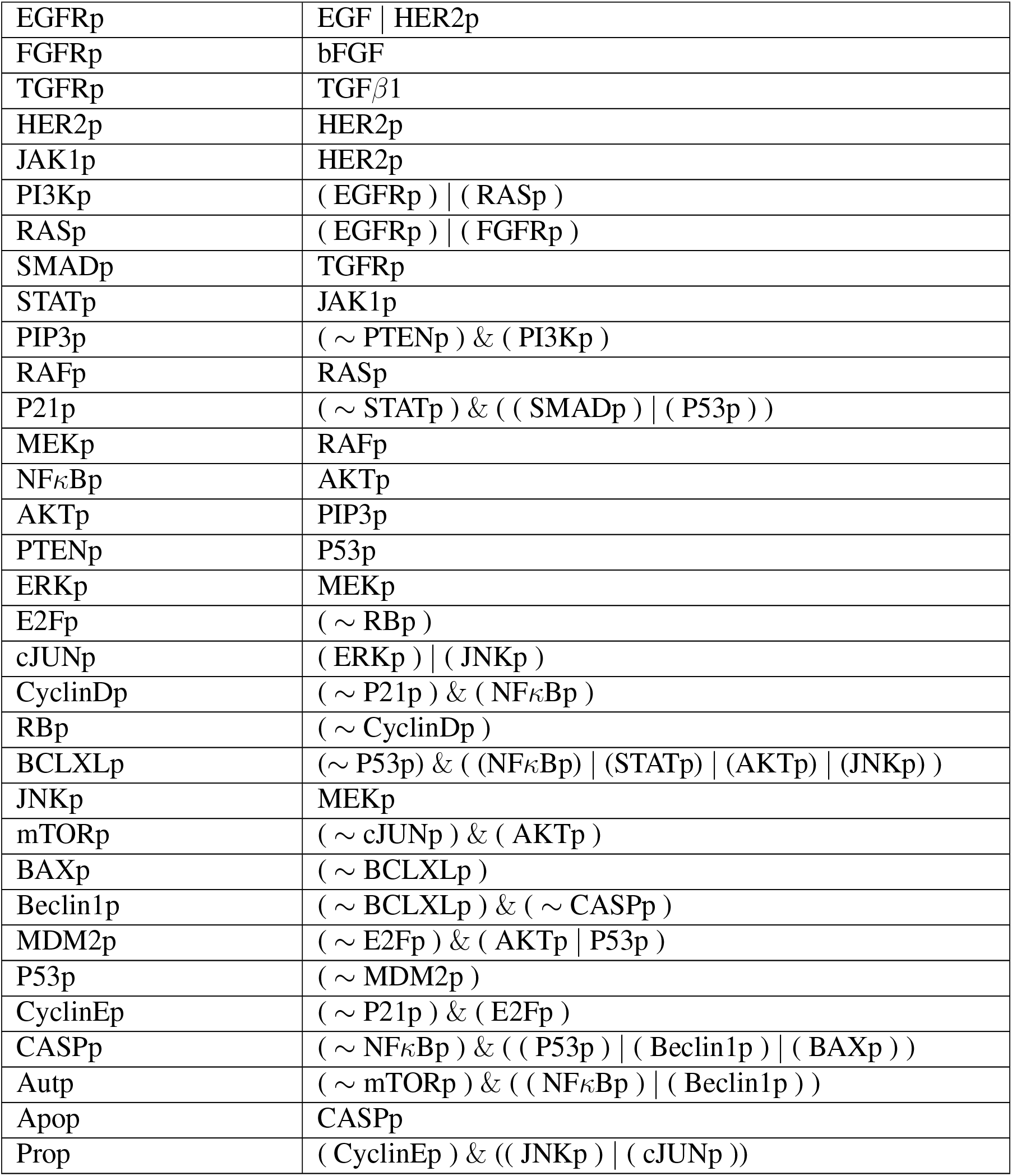
Boolean functions for the whole pancreatic cancer model. Each function indicates the next state of the node in terms of the current states of said nodes’ regulators. Activation is written as OR statements, while suppression is written as AND NOT. The exception to this rule is PCC proliferation, because of its’ upstream signaling.

**Table 6:**
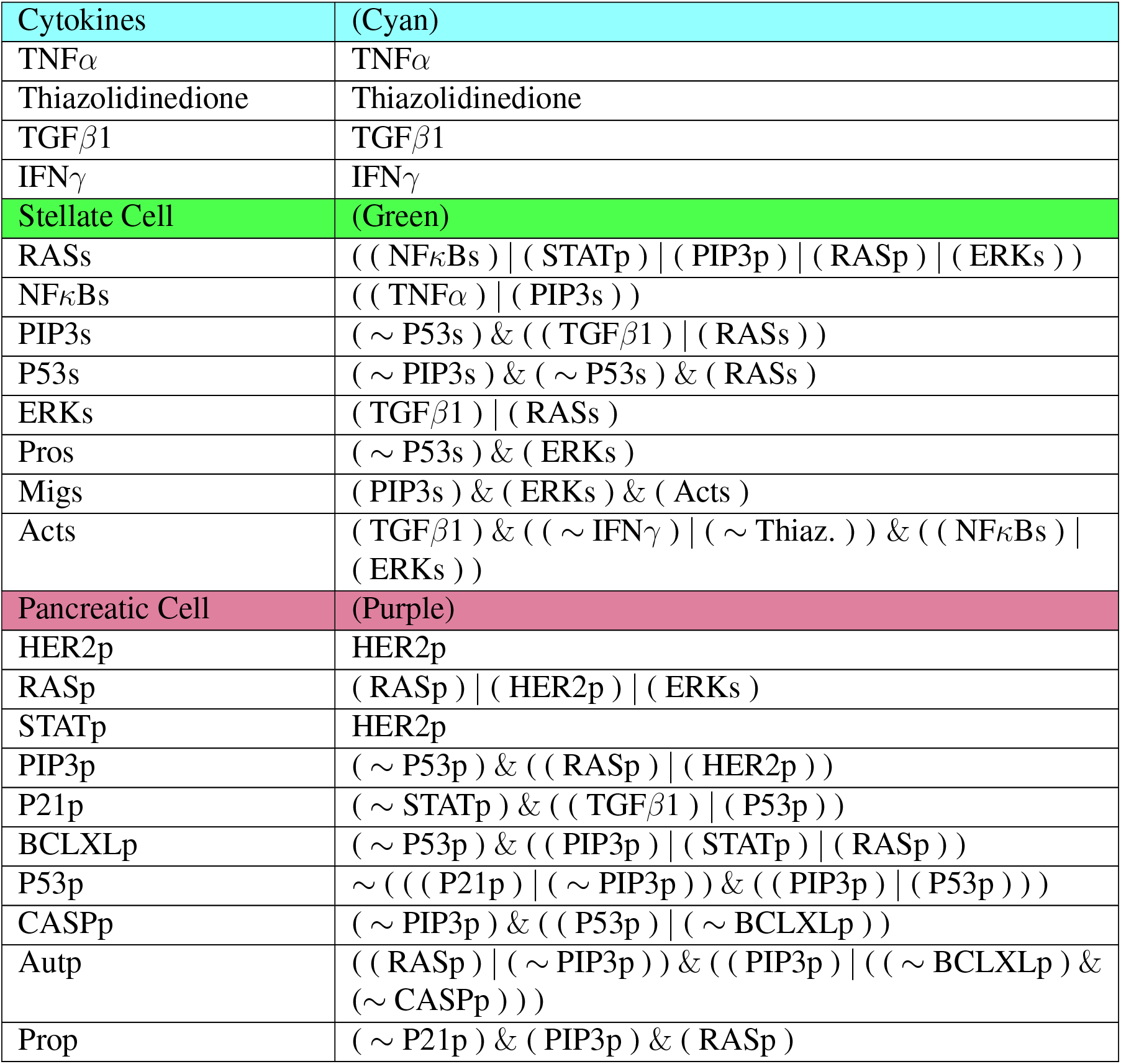
Boolean functions for the reduced pancreatic cancer model. Each function indicates the next state of the node in terms of the current states of said nodes’ regulators. Activation is written as OR statements, while suppression is written as AND NOT. Functions maintain the rules from the whole model by substituting values from the deleted nodes.

### Reduction Examples

For an example of one input and one output, consider FGFRp. The original model’s neighborhood about FGFRp is shown in Figure 4 with equations (1)–(2).

**Figure 4:**
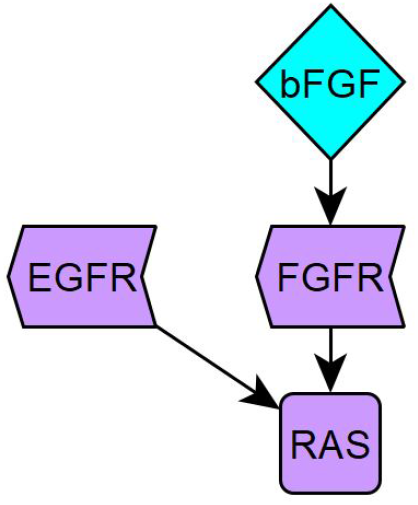
FGFR neighborhood

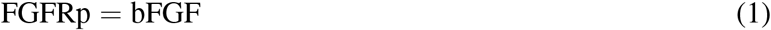

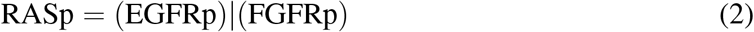

After reduction, we obtain the neighborhood seen in Figure 5 with equations (3)–(4).

**Figure 5:**
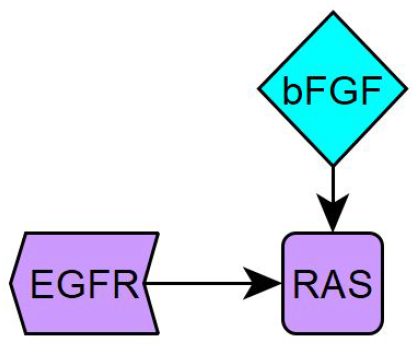
neighborhood around removed FGFRp

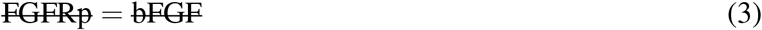

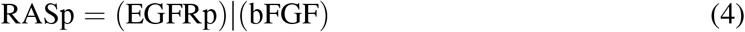

For an example of either one input and multiple outputs, or vice versa, consider MEKp. The original model’s neighborhood about MEKp is shown in Figure 6 with equations (5)–(7).

**Figure 6:**
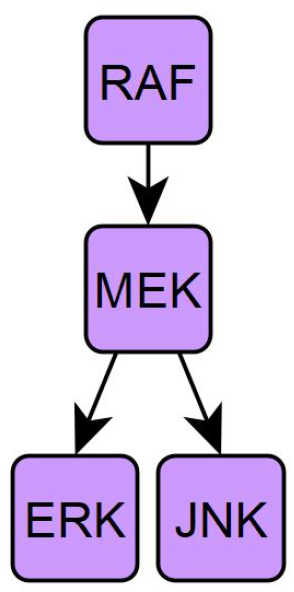
MEKp neighborhood

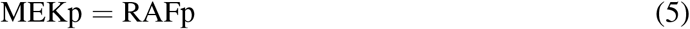

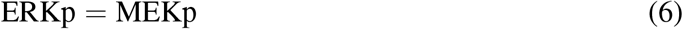

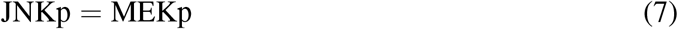

After reduction, we obtain the neighborhood seen in Figure 7 with equations (8)–(10).

**Figure 7:**
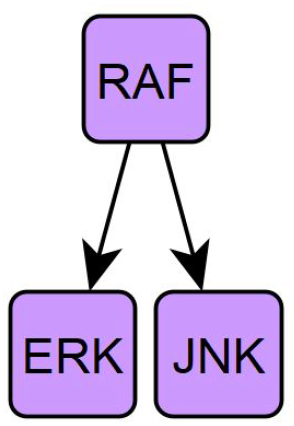
neighborhood around removed MEKp

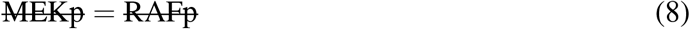

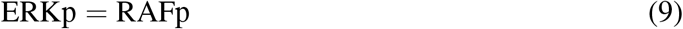

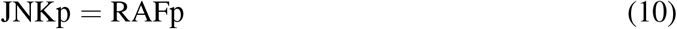

For an example low connectivity, consider cJUNp. The original model’s neighborhood about cJUNp is shown in Figure 8 with equations (11)–(14).

**Figure 8:**
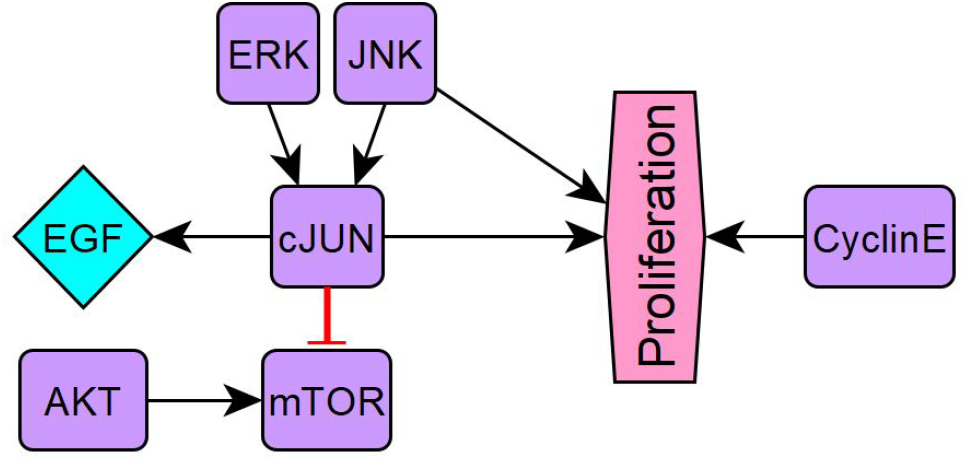
cJUNp neighborhood

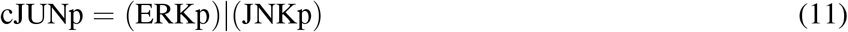

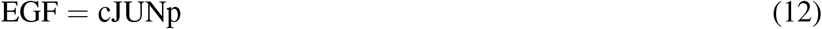

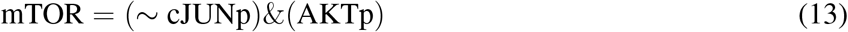

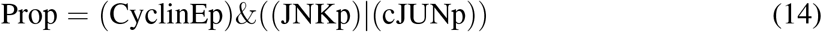

After reduction, we obtain the neighborhood seen in Figure 9 with equations (15)–(18).

**Figure 9:**
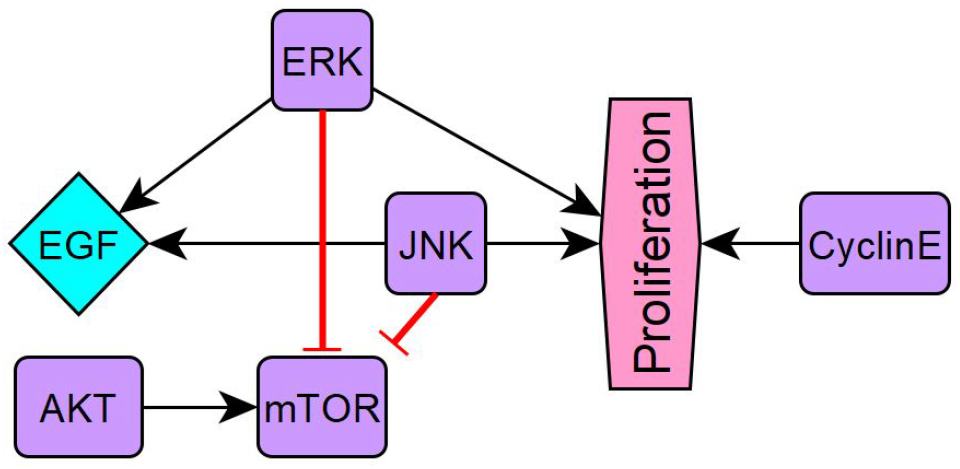
neighborhood around removed cJUNp

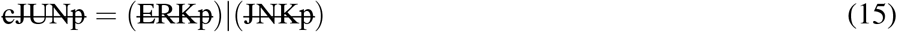

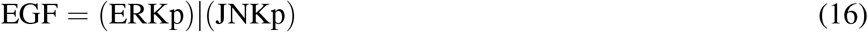

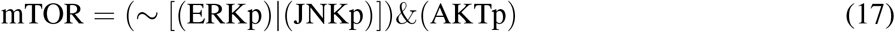

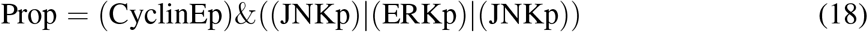

### System Attractors

**Table 7:**
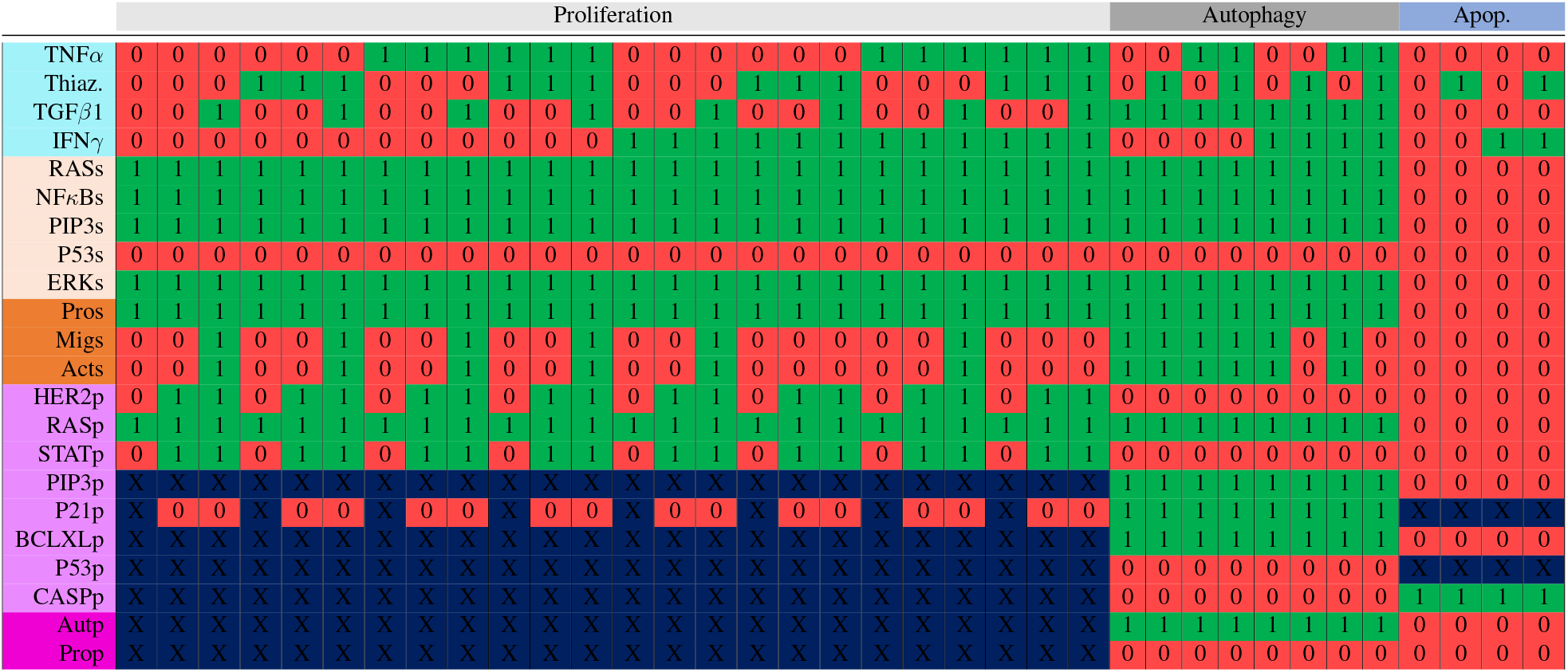
Attractors for the reduced model. We correspond OFF with the Boolean identifier of 0 in red, ON with 1 in green, and cycles are labeled with an *X* in blue. These attractors were found using the stable motif code listed in Section 4. There are thirty-six total attractors, eight of which are fixed points (Autophagy). The reduction left twenty-two nodes, and CASPp identifies PCC apoptosis.

**Table 8:**
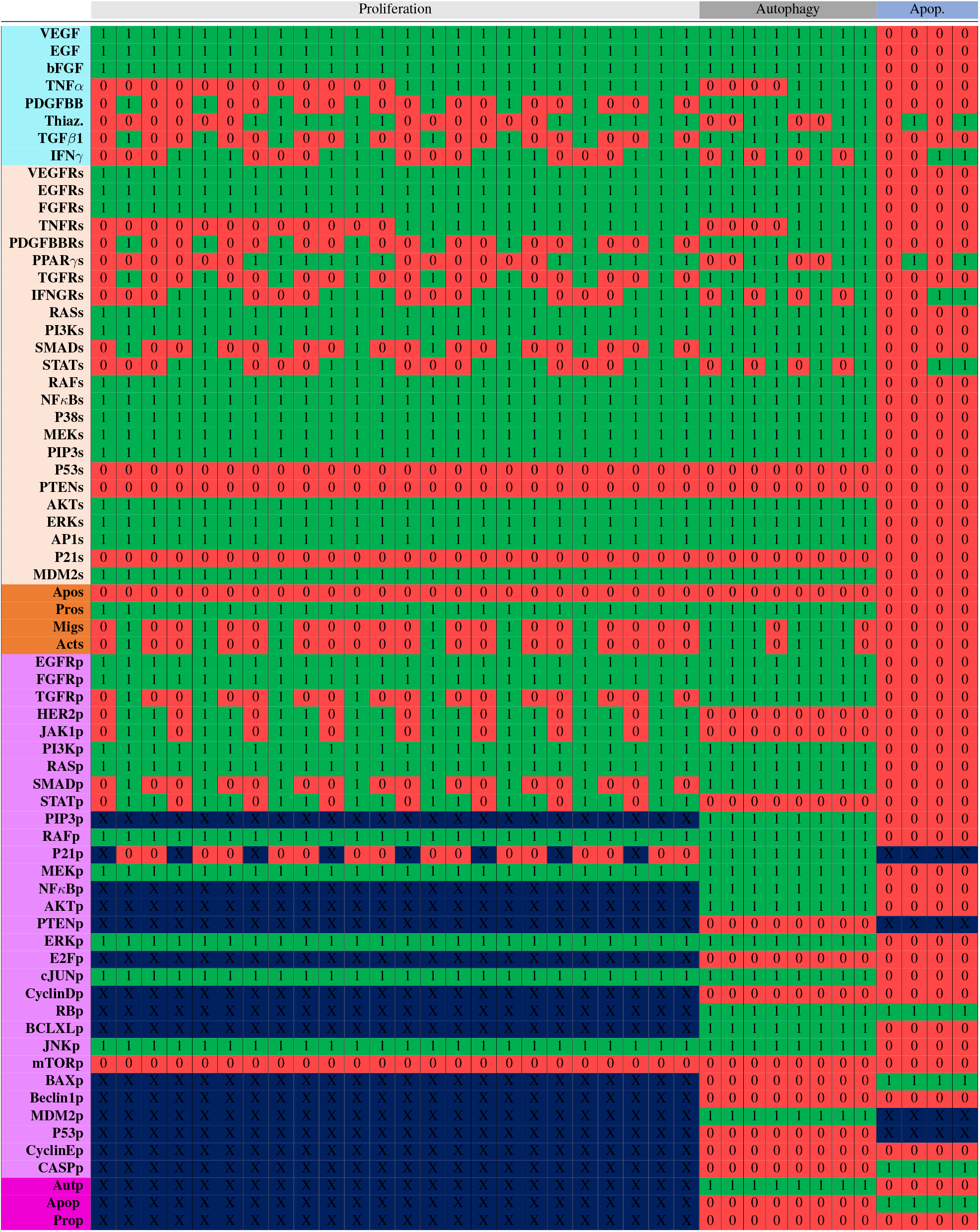
Attractors for the whole model. We correspond OFF with the Boolean identifier of 0 in red, ON with 1 in green, and cycles are labeled with an *X* in blue. These attractors were found using the stable motif code listed in Section 4. There are thirty-six total attractors, eight of which are fixed points (Autophagy).

